# Induction of p73, Δ133p53, Δ160p53, pAKT leads to neuroprotection via DNA repair by 5-LOX inhibition

**DOI:** 10.1101/593202

**Authors:** Shashank Shekhar, Sharmistha Dey

## Abstract

Lipooxygenase-5 (5-LOX), protein is involved in the pathologic phenotype of AD which includes amyloid-plague and tau hyperphosphorylation. This study aims to identify the mechanistic role in neuroprotection by peptide YWCS, the 5-LOX inhibitor in neurotoxic SH-SY5Y cell line developed by the treatment of Aβ_25-35._ The cells were treated with Aβ_25-35_ and with different doses of YWCS. The effect on cell survival pathways were determined by western blot using polyclonal anti body of p53, anti-Akt and anti-phosphorylated-Akt. Immunoprecipitation and mass spectroscopic studies were done to identify the altered proteins. Over expression of phosphorylated-Akt and 3 bands of p53 isoforms were observed which correspond to p73, Δ133p53 and Δ160p53 in the cells treated only with 80µM of YWCS compare to untreated cells. However, no alteration of total p53 and Akt were observed. The results exposed the novel mechanistic pathway of neuroprotection by 5-LOX inhibition, which is likely to be mediated by DNA DSB repair through p53 isoforms and PI3K/Akt pathway. Our finding has opened a new window in the therapeutic approach for the prevention of AD.

## Introduction

Alzhemier’s disease (AD) is the most common cause of dementia in the elderly. Due to increasing longevity and the lack of therapy, AD has become not only a major health problem but also imposes substantial social and economic burden worldwide[1]. The inflammatory process plays a key role in neurodegenerative disorder. The inflammatory molecule, 5-lipooxygenase (5-LOX), protein is involved in the pathologic phenotype of AD which includes Aβ amyloid deposition and tau hyperphosphorylation. Recent research from our lab has proposed the role of 5-LOX peptide inhibitor as neuroprotective molecule for AD[2], provides rescue to neuronal cells from amyloid induced proteotoxic stress/neurotoxicity. The 5-LOX peptide inhibitor reduced γ-secretase expression as well as tau hyperphosphorylation at threonine 181. The mechanistic detail of the 5-LOX inhibitor is still to be elaborated to understand how this inhibitor is providing rescue to neuronal cells from amyloid beta induced neurotoxicity. Some previous studies have shown that 5-LOX inhibitor accelerates the phosphorylation of Akt and thereby provides rescue to cells[3]. It is also reported that 5-LOX regulate p53 activity[4] and p53 isoform Δ113p53/Δ133p53 promotes DNA double-strand break repair to protect cell from death and senescence in response to DNA damage[5]. In a recent finding it has been reported that the p53 isoform; p73 plays an important role in the DNA repair pathway in coordination with another isoform of low molecular weight; Δ133p53[6]. The Δ160p53 is a recently identified isoform of p53 and have role in senescence and DNA repair but the detailed function of it is still illusive[7]. The neuroprotective role for p53 was reported in an in-vivo model of tau-mediated neurodegeneration relevant to Alzheimer’s disease and related disorders. All these findings are illustrated the importance of Akt, p53, p73, Δ133p53 and Δ160p53 in cell survival pathway. Their role in AD pathogenesis is not yet studied. Here, for the first time we have identified the involvement of p73, Δ133p53 and Δ160p53 in amyloid beta induced neurotoxicity and the novel pathway of neuroprotection by YWCS peptide inhibitor of 5-LOX via p73, Δ133p53 and Δ160p53.

## Materials and Methods

### Solid phase Peptide Synthesis and processing

The peptides were synthesized by solid phase peptide synthesizer PS3 (Protein technology, USA) using Fmoc and Wang resin chemistry. The purity of the peptide was verified by analytical RP-HPLC as described earlier[8]. Briefly, the solvent used for the synthesis was dimethylformamide (DMF) and 2-(1H-Benzotriazole-1-yl)-1,1,3,3-tetramethyluronium hexafluorophosphate (HBTU) was used as an activator. Fmoc was deprotected by 20% piperidine and wang resin was cleaved by Trifluoroacetic acid (TFA). The peptides were precipitated from dry ether.

### Preparation of aggregated Aβ peptide

The Aβ_25-35_ peptide was dissolved in nuclease free water and then incubated for 5 days at 37°C and constant shaking in incubator shaker. The aggregation of peptide was confirmed by Thioflavin T (ThT) assay and the scanning electron microscopy.

### Cells and treatment

SH-SY5Y cells were obtained from NCCS, Pune, India and maintained in Ham’s F12 Nutrient media (Gibco) supplemented with 10% (v/v) fetal bovine serum and 1% antibiotic-anti-mycotic solution (Gibco). The cells used in the experiment were of passage number 30-33. Before starting the experiments, cells were authenticated by STR profiling (DNA forensic laboratory Private limited, India). The cells were maintained at 37°C and 5% CO2 under humidified condition. Cells were grown as monolayer.

SH-SY5Y cells were plated at a density of 1×10^6^cells per T25 flask and kept in CO_2_ incubator at 37°C overnight to adhere. Cells were differentiated by the treated of 20 µM retinoic acid in fresh complete media [9,10] for 7 days where media was replaced every 2^nd^ day. On 7^th^ day, cells were treated with different concentrations of YWCS (20, 40 and 80µM) and Aβ_25-35_ (20 µM) peptide for 72 h simultaneously. Aβ_25-35_ and YWCS peptide were synthesized by solid phase peptide synthesis as described previously [8].

### Western blot

Cells were harvested and lysate was prepared in RIPA buffer (10mM Tris-HCl pH-8.0, 140mM NaCl, 1mM EDTA pH 8.0, 0.1% sodium deoxycholate, 1% Triton X100, 0.1% SDS, 0.1mM ethylene glycol tetra-acetic acid pH-8.0, 1mM Protease inhibitor, 1mM phenylmethanesulfonyl fluoride). The expression of p53, Akt and p-Akt were determined by western blotting as described previously [11]. Briefly, 30µg of total protein was separated on 12% SDS gel and protein was transferred on PVDF membrane (mdi, India). Membrane was blocked in 5%NFM and then incubated with primary antibodies of following proteins: p53 (1:500, Santa Cruz), Akt p-Akt, (1:1000, abchem). Membrane were developed with chemiluminescent substrate west Pico (Thermo scientific).

### Immunoprecipitation

After treatment with YWCS cells were lysed in RIPA buffer as described above. Total protein (100µg) was incubated overnight at 4°C with 1µg of anti-human p53 polyclonal antibody in a reaction buffer of 0.1 % BSA in PBS. The protein agarose beads (30µL pre washed with 0.1 % BSA) were added in the reaction and incubated for 4 h at 4°C. After 4 h beads were washed 3 times with wash buffer (10mMTris-HCl pH 7.5, 1mM EDTA, 1mM EGTA, 150mM NaCl, 0.2 mM sodium orthovanadate, 1 mM PMSF). Beads were eluted with 0.2 M glycine pH 2.6 in 1:1 ratio. After elution all the fractions were pulled and neutralized by adding Tris-HCl pH 8.0.

### Mass Spectroscopy

Immunoprecipitated samples were separated on 10 % SDS-PAGE. IgG was used as negative control. Then bands were carefully cut and sent to Sandor Life Sciences, India for the mass spectroscopy.

## Results

### Preparation of aggregated Aβ peptide

Thioflavin-T is a dye specific for detection of fibrillation of proteins. It has an excitation wavelength at 440nm and the emission wavelength at 480nm.ThT assay had shown high intensity after 4 days for it’s aggregation status (Figure 1 A). The results were further confirmed by scanning electron microscopy for the aggregation of peptide after incubation of 4 days (Figure 1 B and C).

**Figure 1.**
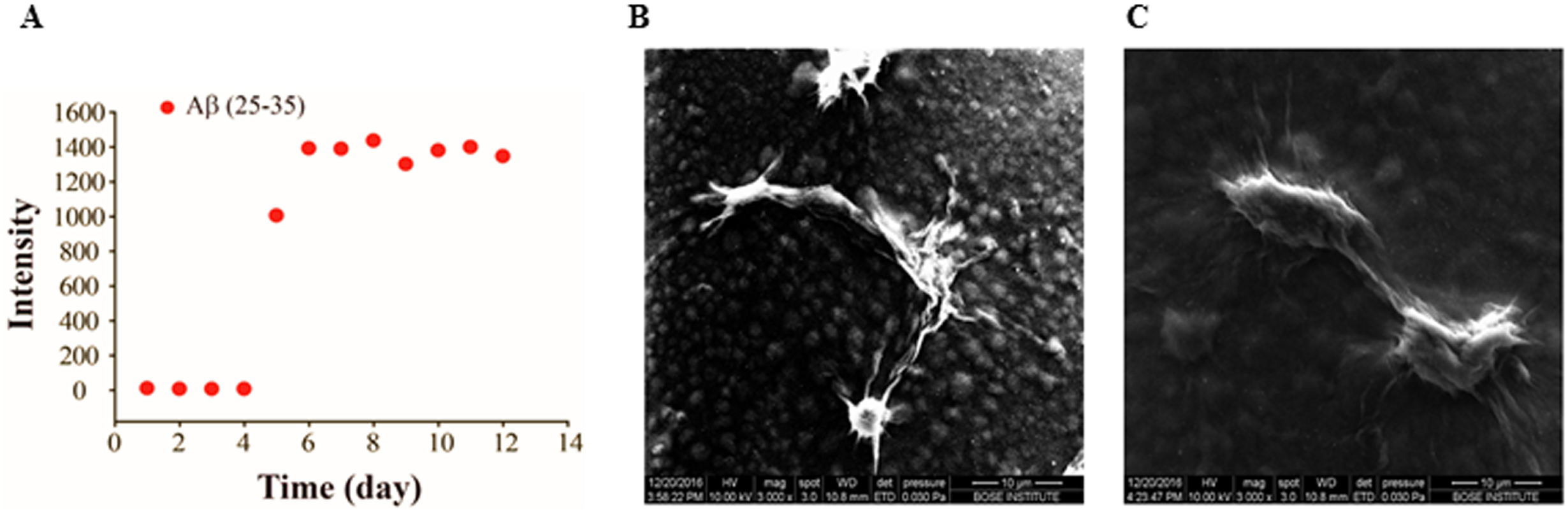
Aggregation of Aβ_25-35_ peptide. (A) There was drastic change in the florescence intensity after 4 days of aggregation. (B) Electron microscopic image of Aβ_25-35_ peptide prior to aggregation. (C) Electron microscopic image of Aβ_25-35_ peptide after aggregation.

### Western blot for Akt and p-Akt

The western blot for Akt and p-Akt were carried out to check the effect of YWCS peptide on their expression and phosphorylation status. Our results suggest that the inhibition of 5-LOX with YWCS peptide had no effect on the Akt expression, while increased the phosphorylated Akt (Figure 2). This indicated that, YWCS peptide induced the phophorylation of Akt protein. Some recent literature reported that the p-Akt has inhibitory effect on apoptosis. Our study first time provides an evidence for phosphorylation of Akt by inhibiting 5-LOX with peptide inhibitor and thereby prevents neurotoxicity.

**Figure 2.**
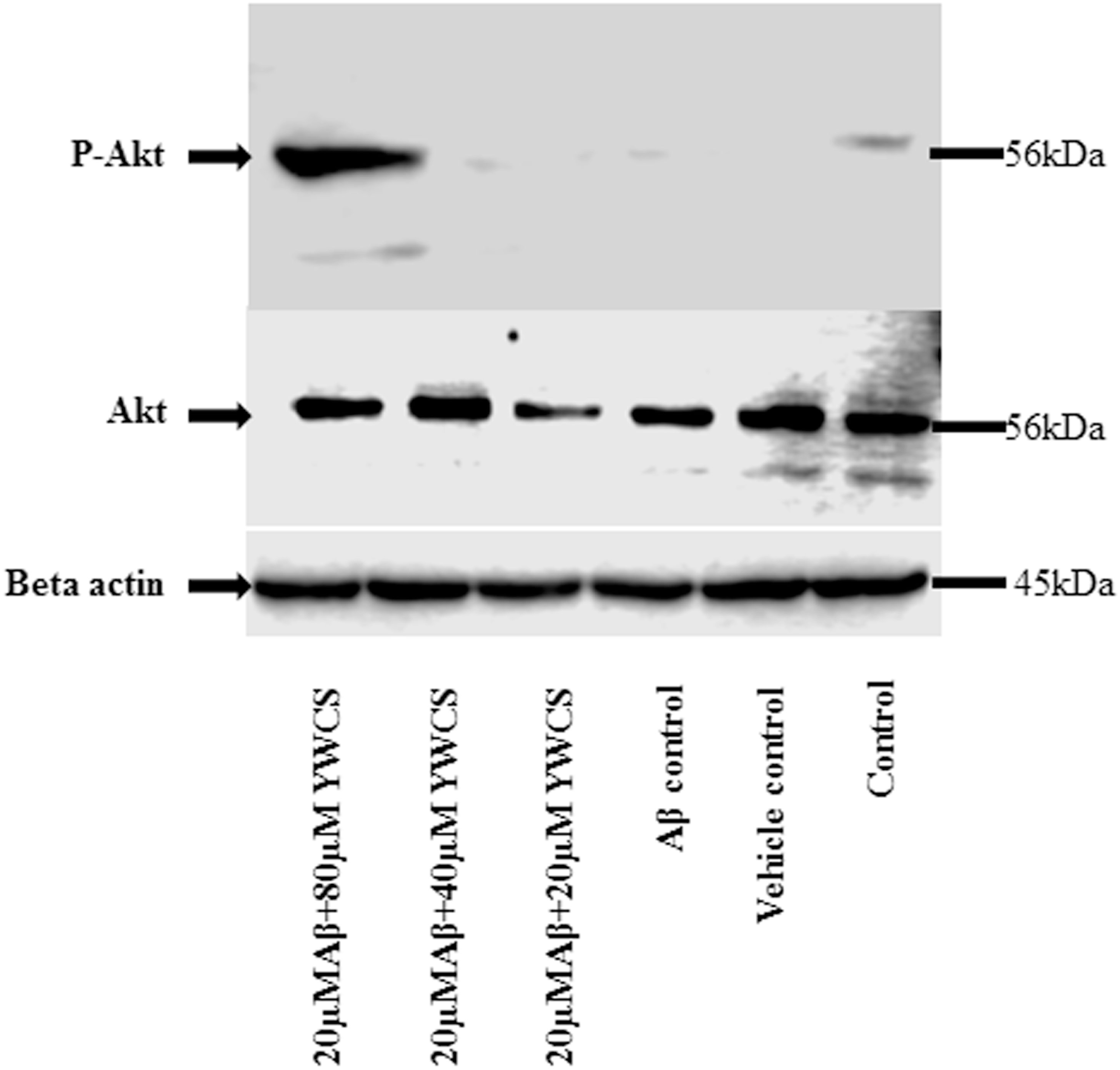
Western blot of expression of Akt and the phosphorylation of Akt at ser473 after treatment by YWCS in SH-SY5Y cells. There was no change in the level of Akt but there was phosphorylation of Akt only in 80 µM of YWCS treated cells.

### Western blot for p53

The western blot for p53 was performed with poly clonal antibody against human p53 since it has several isoforms and play diverse role in cell survival regulation. To our surprise we observed that there was no change in the expression of p53 protein upon treatment with the YWCS but 3 other bands were observed in the cells while treated with 80µM of YWCS. These bands were not present in any other treatment groups (Figure 3).

**Figure 3.**
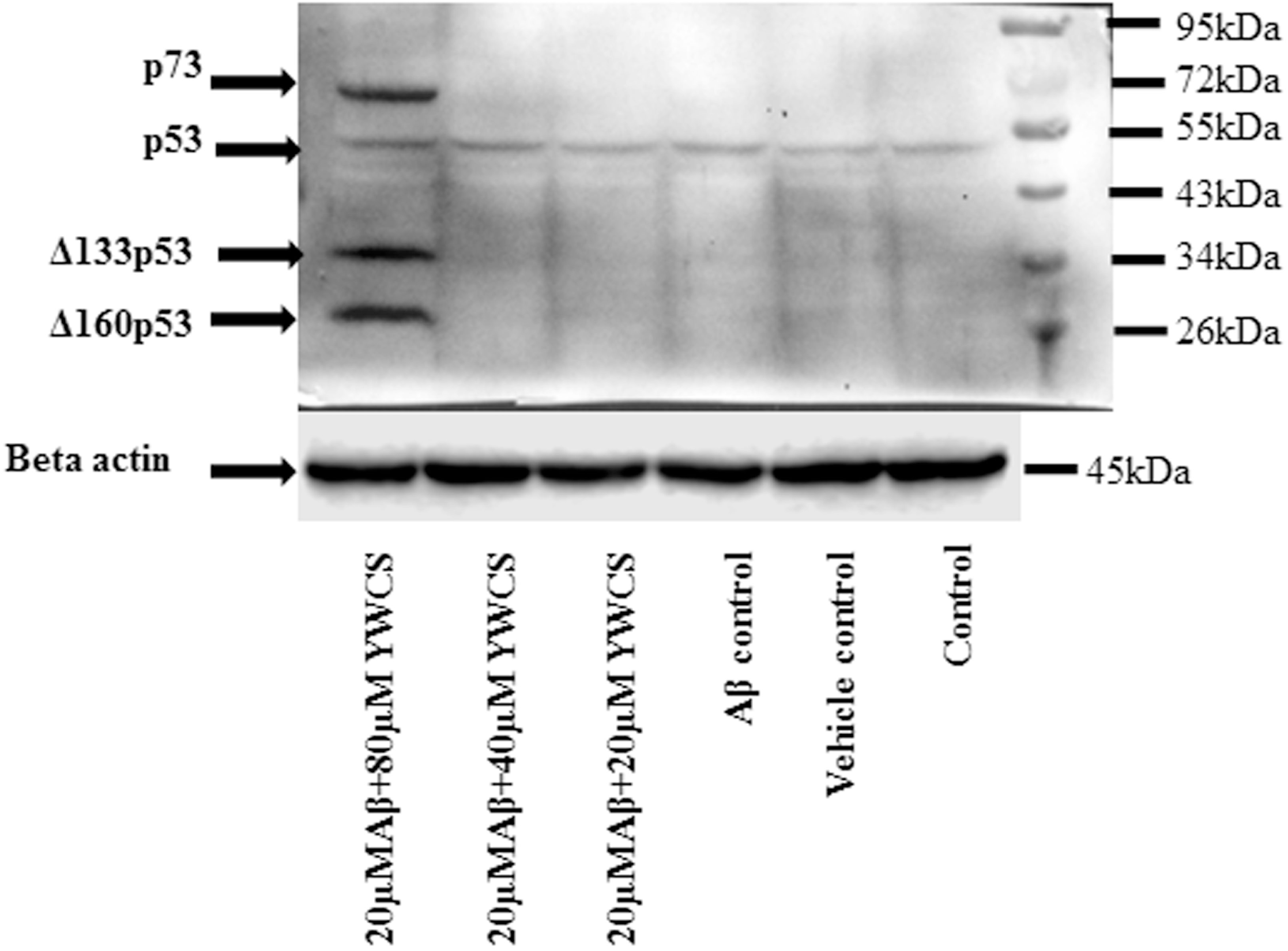
Western blot for p53. Expression of p73 and low molecular weight isoforms of p53 i.e. Δ133p53 and Δ160p53 in 80µM of YWCS treated cells. Beta actin was used as the loading control.

### Identification of proteins which were expressed by the treatment of YWCS

Immunoprecipitation was performed followed by mass spectroscopy. Immunoblot of pull down showed the enrichment of the target proteins (Figure 4A). After immunoprecipitation, pull down was separated on SDS-PAGE and IgG was used as control (Figure 4B). Bands were then sent for mass spectroscopy. The mass result confirmed our hypothesis that the 5-LOX inhibition by peptide induced the expression of p53 isoforms p73, Δ133p53 andΔ160p53 (Figure 5 A, B and C). Our study first time demonstrated the role of p73, Δ133p53 and Δ160p53 in neuroprotection. The expression of these isoforms was induced in the 80µM peptide treated cells. These isoforms of p53 activates DNA double strand break repair. This study showed 5-LOX inhibitor activated these isoforms and prevented neurotoxicity via DNA double strand break repair pathway. This finding opens a new window of mechanistic pathway for 5-LOX inhibition and neuroprotection under oxidative stress and DNA damage.

**Figure 4.**
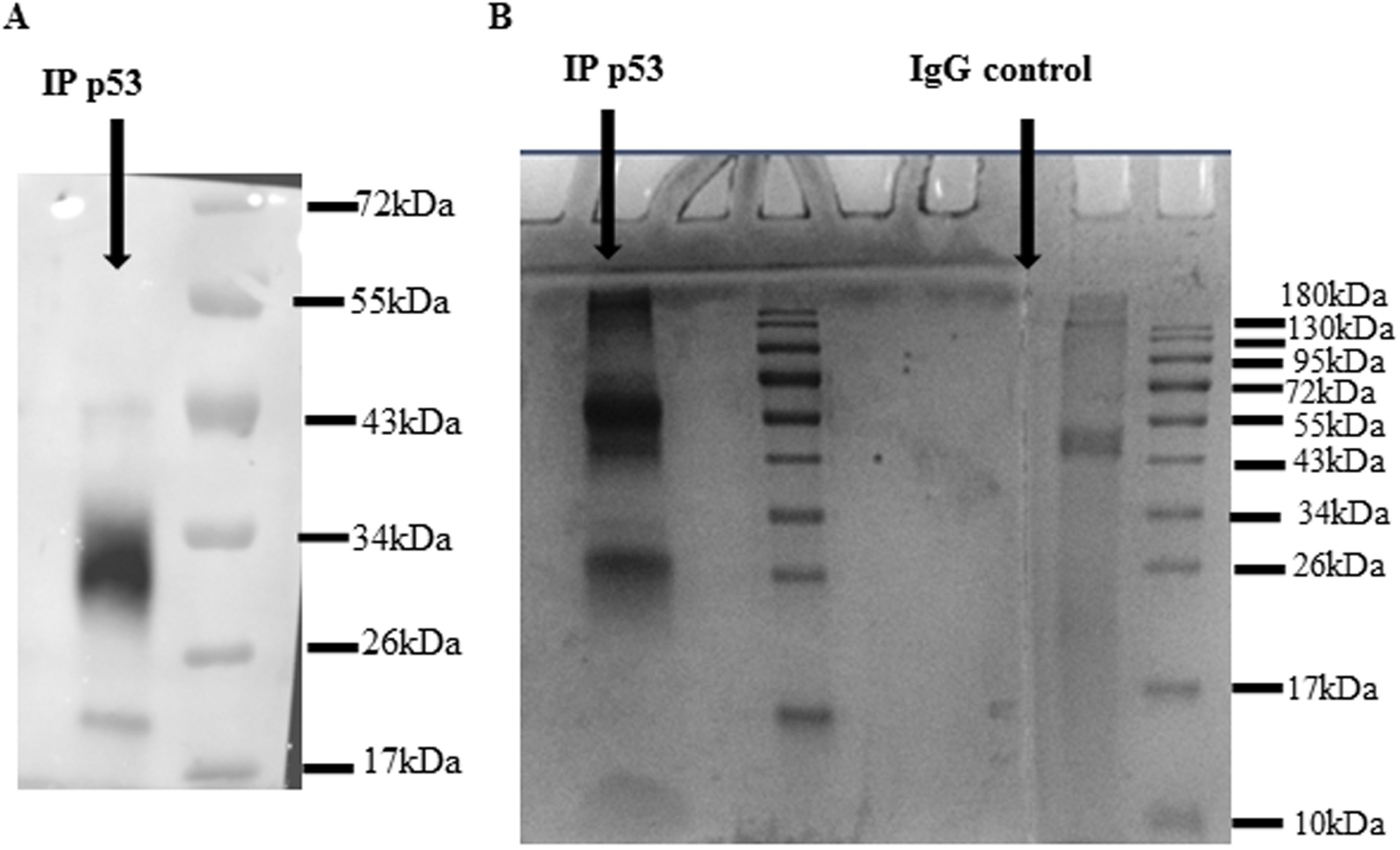
Immunoprecipitation (IP) of p53. (A) immunoblot of IP samples. (B) SDS page of Immunoprecipitate. IgG was used as negative control.

**Figure 5.**
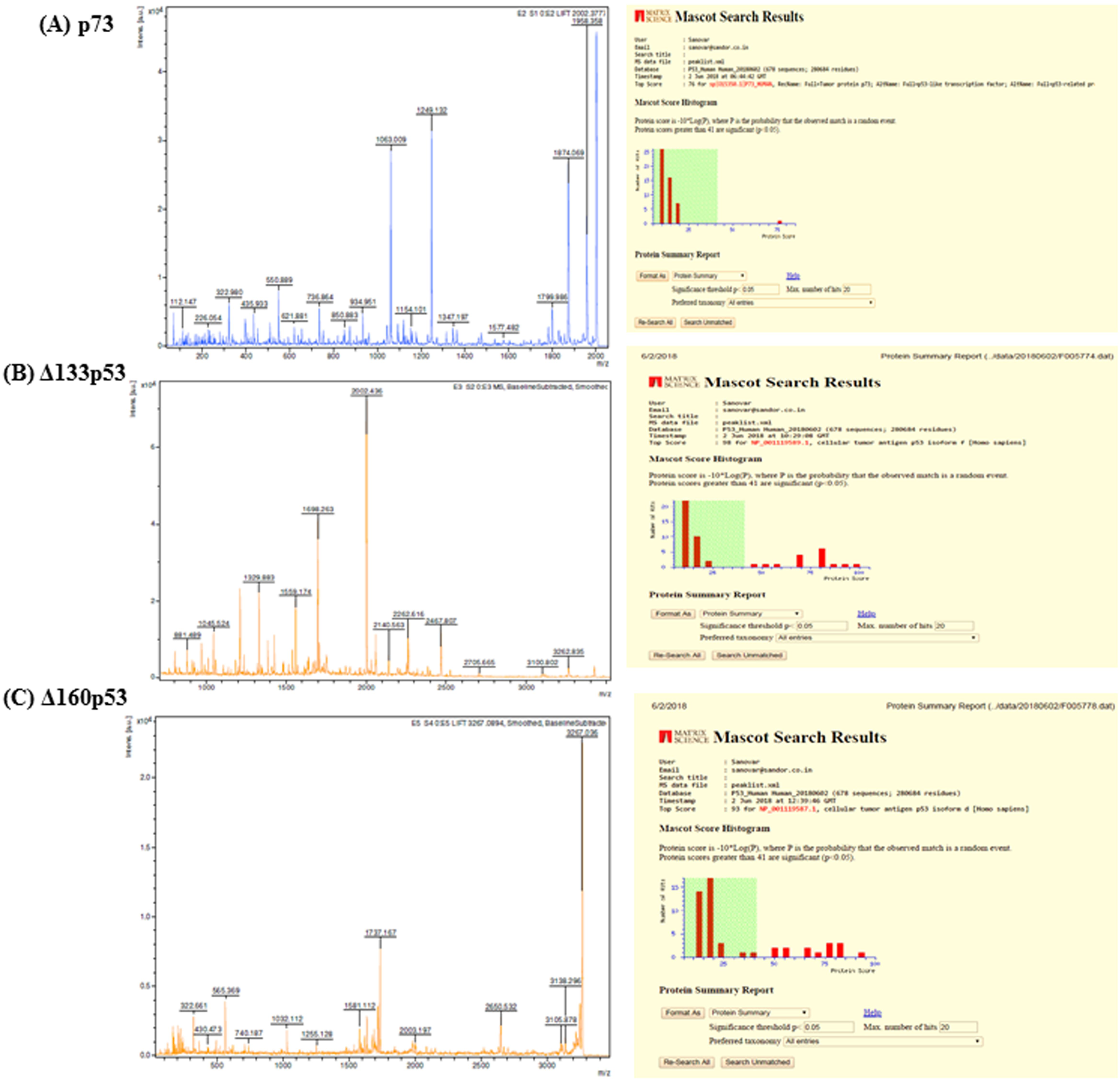
Mass spectroscopy of bands form IP. (A) Identified as p73, (B) identified as Δ133p53 and (C) identified as Δ160p53 isoforms of p53.

## Discussion

AD is age associated disease, progressive with extracellular amyloid-beta (Aβ) deposits intracellular aggregates of hyperphosphorylated tau and neurofibrillary tangles. Ageing process normally arises DNA damage and in AD excessive DNA damage occurs due the Aβ–induced oxidative stress. In our previous work, we found that inhibition of 5-LOX prevented the Aβ induced neurotoxic effect in SH-SY5Y cells and downregulated the expression of γ-secretase components [2]. 5-LOX is a direct p53 target gene in humans. The p53 protein involved in DNA repair, down regulated in AD[12]. In the mammalian cell, p53 was found to be modulate the DNA repair and stimulate both removal of damaged bases and nucleotide re-insertion [13]. To ensure the effect on p53 through 5-LOX inhibition under neurotoxic condition in cells, we performed western blot with treated SH-SY5Y cells by 3 different concentrations (20, 40 and 80 µM) of YWCS using p53 polyclonal antibody against human p53. No alteration of p53 protein was observed, but 4 different bands appeared only at high concentration (80 µM) of peptide. This distinction leads us to search the protein bands by immunoprecipitation and mass spectroscopy, which revealed three isoforms of p53corresponds to p73, Δ133p53 and Δ160p53. While going through literature we came to know that these isoforms play vital and diverse role in cell survival regulation. The isoforms of p53 are found to be pro-survival factor for DNA damage stress and their expression prevents apoptosis and promotes DNA-DSB repair. Some more recent studies have proposed the role of p53 isoforms in the cell survival such as Δ133p53/Δ113p53 by repairing DNA damage in cells during low level of oxidative stress or reactive oxygen species (ROS). However, the full length p53 inhibits DNA-DSB repair. The present work found high expression of p53 isoforms in the Aβ induced neurotoxic SH-SY5Y cells treated by only at higher concentration of peptide YWCS in compare to untreated cells and thereby prevented neurotoxicity. However, no alteration of p53 was observed in treated cells.

The higher expression level of Δ160p53 and p73 isoforms was observed in treated SH-SY5Y cells with peptide (80 µM YWCS). The internal promoter originates the Δ133p53 mRNA codes both the isoforms, Δ133p53 and Δ160 p53. Though Δ160p53 is the conserved isoform, very little information is available about it so far. Another study reported both p73 and Δ133p53 synergistically promote the expression of DNA repairing genes RAD51, L1G4 and RAD52 by homologous recombination (HR), non-homologous end joining (NHEJ) and single-strand annealing (SSA) and thereby promote DNA DSB repairing, which supports our study.

Inhibition of 5-LOX by YWCS peptide inhibitor was also upregulated the phosphorylation of Akt in SH-SY5Y cells. There is evidence of neuroprotective effects by stimulating PI3K/Akt signaling plays a pivotal role in neuronal survival [14]. Previous studies suggested that PI3K/Akt signaling was downregulated in the AD brain [15] and activation of this pathway showed to prevent Aβ-induced neuronal neurotoxicity. Akt has well established role in cell cycle control and in DNA repair by check point activation in late G2 phase.

This study first time reported the rescue of Aβ induced neurotoxicity by the treatment of 5-LOX inhibitor by activating the expression of Δ160p53, p73 and Δ133p53in SH-SY5Y cells. The results reveal that the mechanistic pathway for the prevention of neurotoxic effect by targeting the inhibition of 5-LOX may proceed by DNA DSB repair, by stimulating p53 isoforms and PI3K/Akt pathway. Thus repairing the DNA damage ameliorates the AD pathologies. The results exposed the novel mechanistic pathway of neuroprotection by 5-LOX inhibition mediated by DNA DSB repair through p53 isoforms and PI3K/Akt signaling pathway. Our finding has opened a new window in the therapeutic approach for the prevention of AD.

## Acknowledgments

Authors acknowledge intramural grant, Research Section, All India Institute of Medical Sciences (A-422) for providing funds for the consumable items and CSIR, Government of India for the fellowship of Shashank Shekhar.

## Conflict of Interest

Authors show no conflict of interest.

### Role of the authors

SS performed the entire experiment and study concept. SD was responsible for the study concept, design and wrote the paper.

